# Accurate characterization of CRISPR-Cas9 genome editing outcomes and mosaicism with near-perfect long reads

**DOI:** 10.1101/2025.09.08.674810

**Authors:** Ida Höijer, Robin van Schendel, Anastasia Emmanouilidou, Rebecka Östlund, Ignas Bunikis, Marcel Tijsterman, Marcel den Hoed, Adam Ameur

## Abstract

**Background:** Genetic mosaicism is a consequence of CRISPR-Cas9 genome editing that is difficult to study, especially when it involves structural variants occurring at low frequency. A comprehensive analysis of mosaicism requires deep and unbiased sequencing of the target loci, with accurate single molecule reads. Here we performed amplification-free PureTarget PacBio sequencing to investigate CRISPR-Cas9 outcomes at on-target and off-target sites in germline edited zebrafish and their offspring.

**Results:** Thirty samples from pooled larvae and individual zebrafish were successfully sequenced, resulting in >1100x average target coverage. The PacBio reads reached an exceptional accuracy (QV39) over the target regions, with every read originating from a unique DNA molecule. The two haplotypes of the target loci displayed a balanced depth of coverage, while long-range PCR of the same samples resulted in skewed data. Further analysis of the PureTarget data revealed widespread genetic mosaicism in individual founder (F0) fish, with up to 18 distinct on-target events and 11 off-target events present in a single adult founder. Several CRISPR-Cas9 editing outcomes, including large structural variants and off-target mutations, were inherited to the F1 generation. Notably, as many as seven unique editing events were found among F1 juvenile offspring from a single founder pair, thereby confirming the presence of genetic mosaicism in germ cells of founder zebrafish. This implies that some consequences of CRISPR-Cas9 editing may emerge only in the second generation. We also analyzed DNA methylation but did not observe altered 5mC CpG signals in genome edited samples.

**Conclusions:** PureTarget enables efficient, accurate and unbiased analysis of the full spectrum of genetic mosaicism in a sample. The potential risks of introducing germ cell mosaicism in founder individuals should be carefully evaluated when designing germline genome editing experiments.

## Introduction

Due to its efficiency and ease of use, CRISPR-Cas9 has become an indispensable tool for precise modification of the genome sequence in living cells. In a typical genome editing experiment, the CRISPR-Cas9 complex is directed to its target site by a guide RNA (gRNA) molecule. Once it has reached its target, Cas9 cleaves the DNA molecule with a double-strand break. The broken DNA is then repaired and this process can result in alterations of the DNA sequence at the cleavage site. In absence of a donor template, the DNA repair process often introduces small insertions and deletions, but to some extent also larger structural variants (SVs), and possibly even complex events such as whole chromosome deletions or chromothripsis(1–4). Occasionally, the gRNA can bind to other locations than the intended (on-target) site, leading to off-target genome editing(5,6). Since it may be difficult to foresee the exact outcomes of a CRISPR-Cas9 experiment, the on-target and off-target regions are often verified by DNA sequencing to rule out the presence of undesired genome editing at the on-target and off-target sites.

CRISPR-Cas9 can give rise to multiple editing outcomes at a target site(7–9). This means that a genome edited sample can contain a mosaic of small and large genetic variants, each of which could be either common or rare, and located at the on-target or at an off-target site. This CRISPR-induced mosaicism can be found both in cell samples and in living organisms where different edits may be present in different cell types or tissues. Genetic mosaicism has been found also in founder animals resulting from germline editing of fertilized eggs or embryos, and it is believed that the rate of mosaicism can be accelerated by the CRISPR-Cas9 complex remaining active through several cell divisions(10). A recent study employing targeted amplicon sequencing in single cells revealed a unique editing pattern in nearly every edited cell(11). However, the full extent of CRISPR-Cas9 induced genetic mosaicism occurring in a tissue sample, or in an organism, has not yet been possible to determine. New and improved sequencing methods are required to remove technical hurdles so this phenomenon can be studied in more detail.

The ideal sequencing approach for in-depth analysis of CRISPR-induced mosaicism needs to fulfill some key criteria. Firstly, the sequencing reads must be long enough so that not only small variants but also many of the larger SVs can be detected. Secondly, the data needs to have high quality so the exact variant breakpoints can be determined in individual reads. Thirdly, the method needs to yield high-coverage unique molecule data over the targets. For detection of low frequency variants down to 1% at least 200x coverage is required. Lastly, the sequencing data must be unbiased so that the identified variants represent the true distribution of editing outcomes occurring in the sample. The three first criteria can be met by long-range PCR (LR-PCR) and subsequent long-read sequencing, a method that can be further refined by unique molecular identifiers (UMIs) to remove duplicate molecules and increase the accuracy(12,13). However, even with UMIs, PCR amplification has intrinsic biases, particularly when amplifying longer regions. An amplification-free sequencing approach is therefore preferable.

Amplification-free sequencing can be achieved using instrumentation from Oxford Nanopore Technologies (ONT) or Pacific Biosciences (PacBio)(14). These long-read sequencing technologies not only provide DNA sequence information but also signals for base modifications like CpG 5mC methylation(15,16). Whole genome long-read sequencing (WG-LRS) gives an unbiased and complete representation of the genome. However, it is not suitable for studying mosaicism since the cost of generating >200X coverage WG-LRS data is prohibitively high when investigating many samples. Furthermore, deep WG-LRS requires substantial amounts of DNA that may be difficult to obtain from genome edited samples. A more feasible option is ONT’s adaptive sampling, where target molecules are selected during sequencing(17,18). Adaptive sampling has been seen to generate 9 to 40x median target coverage, or 4.6-fold enrichment when run on a MinION flow cell(19). This means that at most a handful of samples can be multiplexed on a PromethION flow cell in order to reach sufficient target coverage for mosaicism analysis, an experimental setup that still would be relatively costly. Furthermore, ONT sequencing has limitations in indel calling(20), potentially rendering it suboptimal for analysis of mosaicism in CRISPR-Cas9 edited samples, which often consist of small insertions and deletions.

An alternative approach capable of generating deeper coverage over the targets is Cas9-based enrichment in combination with long-read sequencing(21). Such protocols have been developed both for ONT and PacBio, primarily with the aim of resolving repeat expansions in neurological disorders(22,23). ONT’s Cas9 enrichment has been applied for genome editing verification with promising results, although the error rate limits the usefulness of the method and complicates the analysis(24,25). PacBio recently released the PureTarget Cas9-enrichment method along with a panel of gRNAs to target 38 disease causing repeat loci in the human genome. The workflow allows multiplexing of up to 48 samples on a single Revio SMRTcell. In PacBio sequencing, the molecules are circularized by ligating hairpin adaptors, and each base is sequenced several times to generate very high accuracy (HiFi) data over the target regions(26). With PureTarget, the enriched DNA is shorter than the 15-20 kb length in a typical PacBio HiFi library, thereby enabling additional passes of the circular template during sequencing, which could further increase the accuracy. Thus, we reasoned that PureTarget, due to its specific properties, might be able to generate single-molecule reads with sufficient target coverage and quality for detailed investigation of genetic mosaicism in CRISPR-Cas9 genome edited samples.

Here we applied PureTarget to study genetic outcomes of CRISPR-Cas9 editing. The samples were previously examined by us using LR-PCR as part of a study published in 2022, where off-target genome editing and large structural variant events were studied in two generations of zebrafish(4). In the present study, we revisit the zebrafish DNA samples and evaluate PureTarget for simultaneous validation of genetic and epigenetic (5mC) variation induced by CRISPR-Cas9, with the aim to investigate mosaicism at an unprecedented level of detail.

## Results

### High-quality long-read sequencing of CRISPR-Cas9 genome editing outcomes using PureTarget

PureTarget is an amplification-free protocol for targeted PacBio sequencing, initially developed for re-sequencing of repeat expansions in human genomes. The method is based on Cas9 enrichment of regions of interests (ROIs). Here, we adapted PureTarget to investigate genome editing outcomes (Figure 1A) and applied it to DNA samples from two generations of CRISPR-Cas9 germline edited zebrafish previously analyzed by long-range PCR (LR-PCR) and PacBio sequencing(4) (Figure 1B). Four on-target genome editing sites were investigated in the zebrafish samples, targeting coding sequence in the genes *ldrla, nbeal2, sh2b3* and *ywhaqa*. Three previously identified off-target sites were examined as well, one for *sh2b3* and two for *ywhaqa* (Figure 1C and supplementary Table S1). The ROIs were designed to have a length of approximately 5 kb. The 32 samples consisted of six pools of 25-30 10-day-old edited founder larvae, six pools of 30 5-day-old edited F1 larvae, ten individual edited adult founder fish, nine individual edited juvenile F1 fish, and one pool of unedited founder larvae serving as a control sample (supplementary Table S2). The input samples were partially degraded (supplementary Figure S1) due to difficulties to extract intact DNA from zebrafish. Despite this, PacBio sequencing of the 32 PureTarget samples on a single Revio 8M SMRTcell generated 1.8 million reads (supplementary Table S3). 17.1% of reads were aligned to the ROIs, corresponding to an average enrichment rate of 4500x for target regions vs. the rest of the zebrafish genome (supplementary Table S4 and Methods). Target PacBio HiFi reads have an average QV score of 39 (supplementary Figure S2). Although not an exact measure of quality, QV39 corresponds to one error in 7900 bases, implying that many reads mapping to ROIs are completely error-free. The PureTarget assay thus enables single molecule sequencing of the CRISPR-Cas9 editing sites with near-perfect kilobase-length reads.

**Figure 1:**
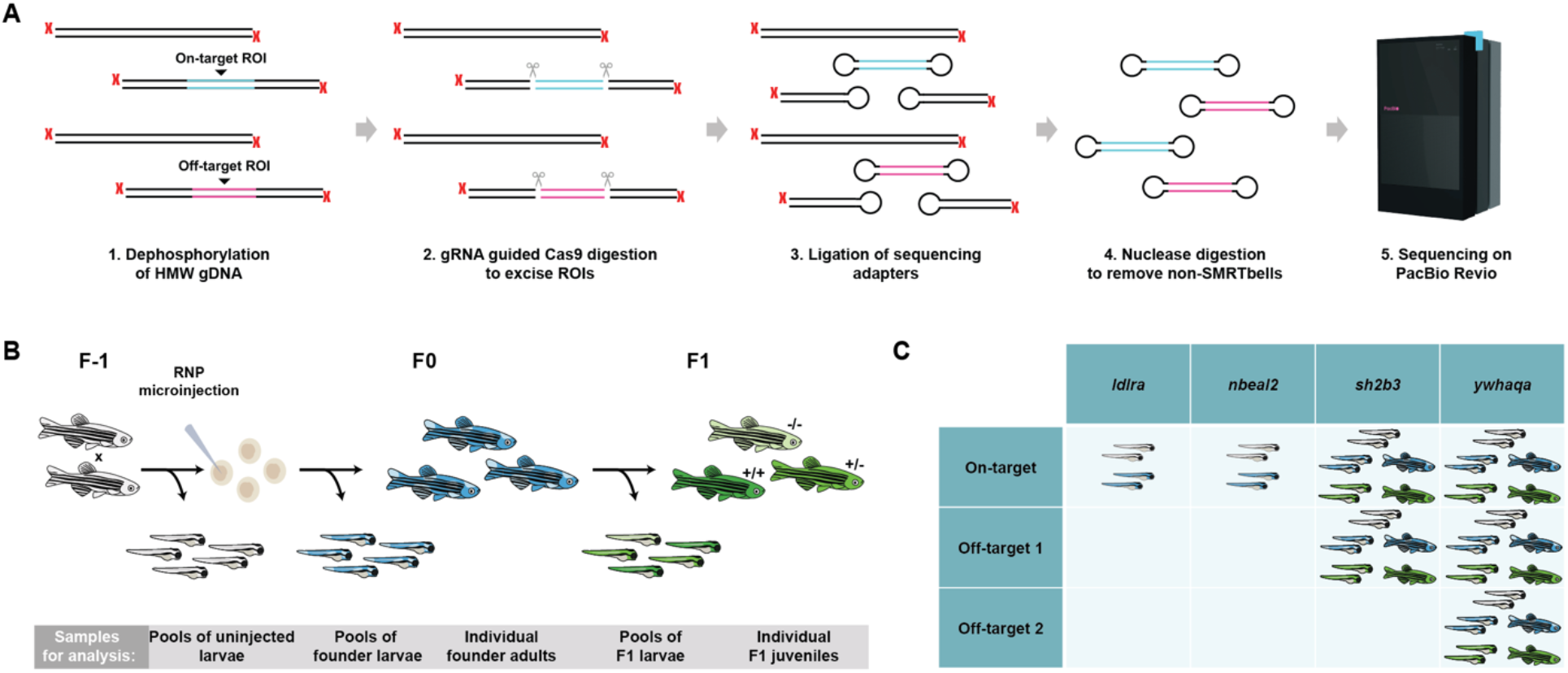
Overview of the targeted sequencing method and samples used in the study. **A)** The steps of the PureTarget protocol for Cas9-based target enrichment and PacBio sequencing. **B)** Overview of the zebrafish experiment. CRISPR-Cas9 genome editing was performed by micro-injection in fertilized eggs at the single cell stage. Samples were obtained from pooled larvae and individual zebrafish from the founder (F0) and F1 generations. **C)** Four gRNAs previously shown to induce off-target mutagenesis were used for CRISPR-Cas9 gene editing of ldrla, nbeal2, sh2b3 and ywhaqa. The table shows the fish samples that were assayed at the on- and off-target sites for the four different gRNAs. In the table, the larvae and fish are color-coded according to the experimental outline in panel B with unedited samples (in gray), edited F0 samples (blue) and edited F1 samples (green).

### Unbiased and in-depth analysis of on-target and off-target editing sites in zebrafish

Two samples from adult founder fish generated low (<30x) coverage of the targeted regions, because of suboptimal Cas9 enrichment or poor DNA quality. These two samples were excluded from the downstream analysis. The 30 remaining samples reached 1168x coverage on average at the seven ROIs, with some samples having >3000x coverage at specific targets (Figure 2A). We were further interested to evaluate the coverage uniformity for the two haplotypes in individual zebrafish. For this purpose, we analyzed the allelic distribution at heterozygous SNVs in individual zebrafish from the F1 generation. We did not include founder fish in this analysis because they may have a skewed allele frequency due to loss of heterozygosity and mosaicism induced by CRISPR-Cas9 editing. In total, eleven heterozygous on-target and off-target regions were found in seven individual F1 fish. A representative SNV position was selected in each target as a measure of allelic distribution. We then compared the allele frequencies at the SNV position in the PureTarget results to LR-PCR data from the same samples (supplementary Table S5). Our results show that PureTarget gives a near 50-50% distribution between the reference and alternative alleles, also in cases where LR-PCR is highly biased (Figure 2B). Viewing the data in IGV(27) demonstrates the advantages of PureTarget when it comes to generating high-quality reads (Figure 2C) and equal representation of the two haplotypes (Figure 2D). The exceptional quality of the PureTarget reads alongside its unbiased nature makes the method ideal to investigate low-frequency and mosaic events in genome edited samples.

**Figure 2.**
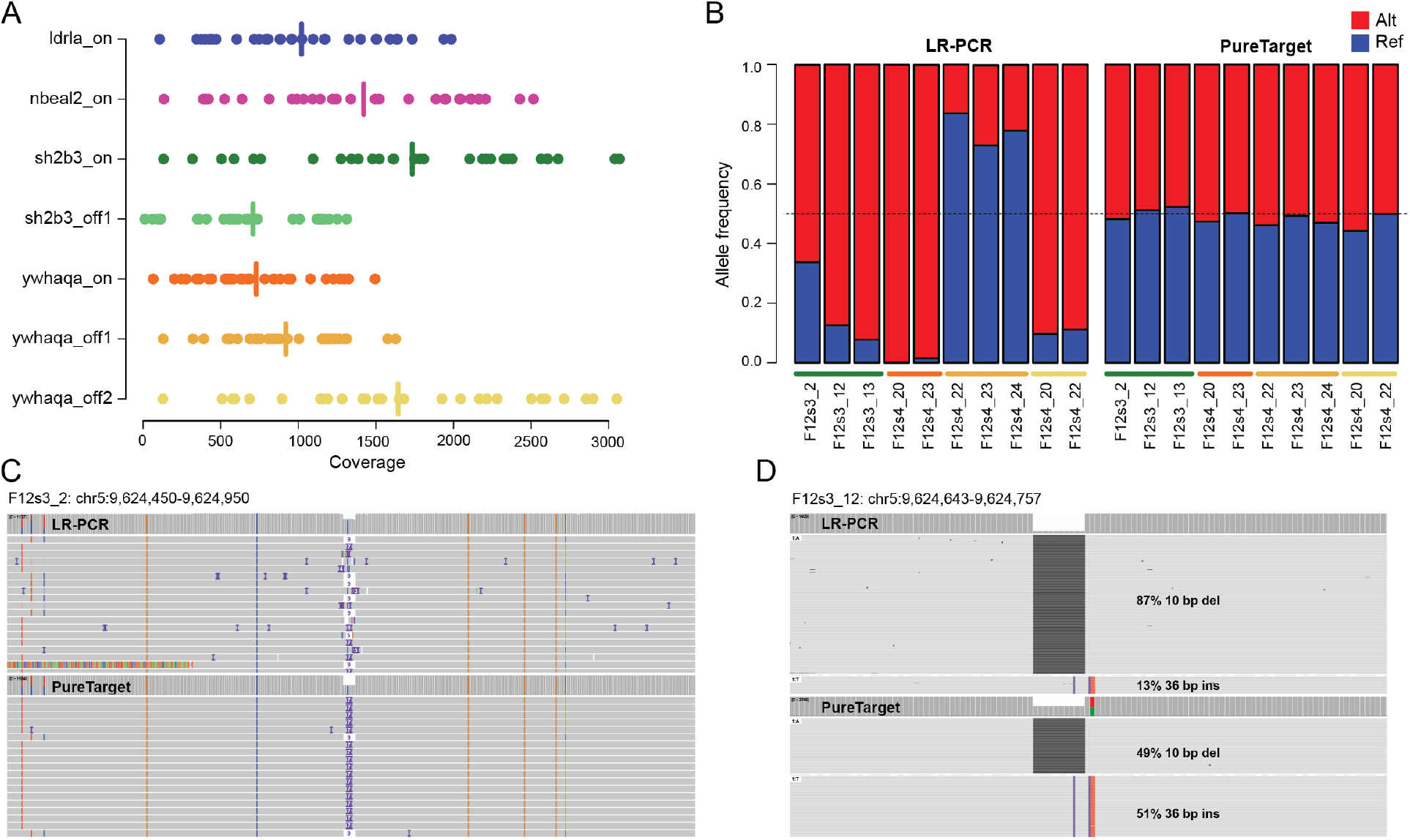
Performance of the PureTarget method on genome edited zebrafish samples. **A)** Coverage across the 30 PureTarget zebrafish samples and seven target regions included in the assay. The vertical lines correspond to the average coverage for the different targets. **B)** Frequencies of the reference (blue) and alternative (red) alleles at single nucleotide variant (SNV) positions in heterozygous target regions for F1 zebrafish. The bars to the left show allele frequencies for long-range PCR (LR-PCR) while the bars to the right show allele frequencies for PureTarget in the same samples. The minor allele frequencies obtained by PureTarget are close to 50% while LR-PCR results in more uneven distribution between the two haplotypes. The colors below the bars indicate which of the target sites from A were examined. **C)** IGV visualization of the sh2b3 on-target region in an F1 juvenile fish, centered around the CRISPR-Cas9 genome editing site. The IGV tracks show representative PacBio read alignments for LR-PCR (top) and PureTarget (bottom), although not all reads are displayed in this view. The PureTarget data contains fewer alignment errors and displays a clearer view of the editing outcomes. **D)** Visualization of the sh2b3 on-target region in an F1 juvenile fish showing results for LR-PCR (top) and PureTarget (bottom). The view has been compressed so all reads are shown for each sample. The sample contains two CRISPR-Cas9 editing outcomes, a 10 bp deletion and a 36 bp insertion. In the LR-PCR data the 10 bp deletion is found in 87% of the reads. For PureTarget, the two alleles have an even balance with 49% of the reads containing the deletion and 51% containing the insertion.

### Founder zebrafish are mosaic at on- and off-target CRISPR-Cas9 editing sites

Having verified the sensitivity of our method for targeted sequencing of zebrafish DNA, we were interested to investigate the level of mosaicism occurring in the founder (F0) generation. For this purpose, we analyzed the PureTarget data using the software SIQ for analysis and SIQplotteR for visualization(28). We compared the mutational outcomes in samples of pooled founder larvae analyzed both by PureTarget and LR-PCR. This revealed a wide range of editing events at the on-target sites, while – as expected – no editing was seen in the control sample (Figure 3A and supplementary Figure S3). Editing was also observed at off-target sites, with the highest degree occurring at *ywhaqa* off-target 2 (Figure 3B and supplementary Figure S4). In the samples of pooled founder larvae there was a large overlap between the on-target events identified by PureTarget and LR-PCR, although the allele frequency distributions showed a certain degree of variability (Figure 3C). The discrepancies in allele frequencies are likely caused by biases in the LR-PCR data. We next examined the PureTarget data for individual adult founder zebrafish. The SIQ results revealed a high degree of on-target mosaicism for *sh2b3* and *ywhaqa*, with between 7 and 18 distinct events present in the founder individuals (Figure 3D). These numbers were obtained by only considering events supported by at least 1% of the reads. Given its high sensitivity, PureTarget likely allows detection of variants occurring at even lower frequencies. As expected, only the wild-type allele reached the 1% threshold for the control sample. Mosaicism was identified also at off-target sites, especially for *ywhaqa* off-target 2, where 9-11 distinct alleles were seen in the adult individuals (Figure 3E). Based on these results, we conclude that genetic mosaicism is highly abundant in adult founder zebrafish.

**Figure 3.**
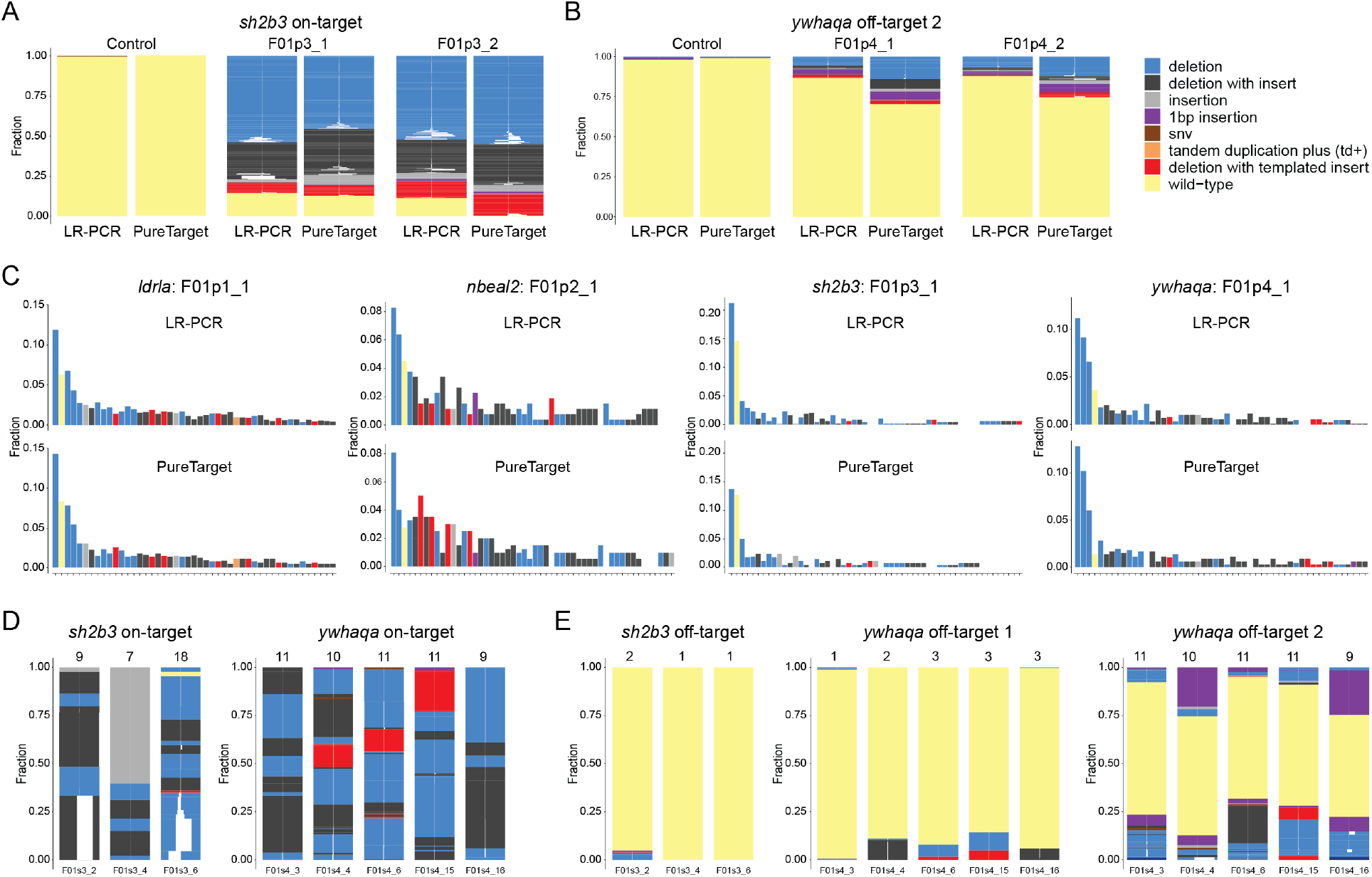
CRISPR-Cas9 induced genetic mosaicism in founder zebrafish. **A)** SIQ tornado plots showing the distribution of on-target genome editing outcomes in the uninjected control sample and two pools of sh2b3 founder (F0) larvae. For each sample, outcomes are shown both for LR-PCR and PureTarget data in a ±2 kb window centered around the CRISPR-Cas9 editing site. The pools of larvae contain many distinct genome editing outcomes while only the wild-type allele is detected in the control sample. **B)** SIQ tornado plots for uninjected control sample and two pools of ywhaqa founder (F0) larvae at ywhaqa off-target site 2 (1 kb window size). **C)** Comparison of CRISPR-Cas9 genome editing outcome frequencies in ldrla, nbeal2, sh2b3 and ywhaqa in pooled founder larvae. For each sample of pooled founder larvae a comparison is shown between LR-PCR (top) and PureTarget (bottom). Each plot shows the ∼50 highest frequency editing outcomes. There is a good overlap between events detected by LR-PCR and PureTarget, but with varying frequencies for some events. **D)** SIQ visualization of all editing events detected at sh2b3 and ywhaqa on-targets in adult founder individuals. At the top of each bar are the number of distinct events with allele frequency >1%. **E)** Editing events at off-target sites for the same founder individuals as in D.

### CRISPR-Cas9 induced mosaicism is present also in germ cells of founder zebrafish

We observed a wide range of genome-editing events in the founder zebrafish and examined how many of these were passed on to the next generation. In our experiment, we sequenced six samples of 30 pooled F1 larvae as well as nine samples from fin-clips of individual juvenile F1 zebrafish. The samples were collected from the offspring of two founder pair crossings, one for *sh2b3* and one for *ywhaqa*. From the offspring of the *sh2b3-*edited founder pair we generated PureTarget data for three F1 larvae pools and for five juvenile F1 fish. Three F1 larvae pools were sequenced also from the *ywhaqa*-edited founder pair offspring, and four juvenile F1 fish. For *sh2b3*, seven on-target editing events were present in all three samples of pooled larvae and in ≥1 of the juveniles (Figure 4A). For *ywhaqa*, five on-target events were overlapping between all three larvae pools and at least one juvenile F1 fish (Figure 4B). For the two investigated founder pairs, we thus found 5 and 7 unique on-target CRISPR-Cas9 editing outcomes among their juvenile F1 offspring. This observation can only be explained by the presence of mosaicism in the germ cells of the founder zebrafish. Several of the events detected in multiple pools of larvae were not seen in the juvenile F1 zebrafish. The absence of these events could be explained by the low sample size of F1 juveniles in our experiment and/or by a selective disadvantage of certain mutations during zebrafish development.

**Figure 4.**
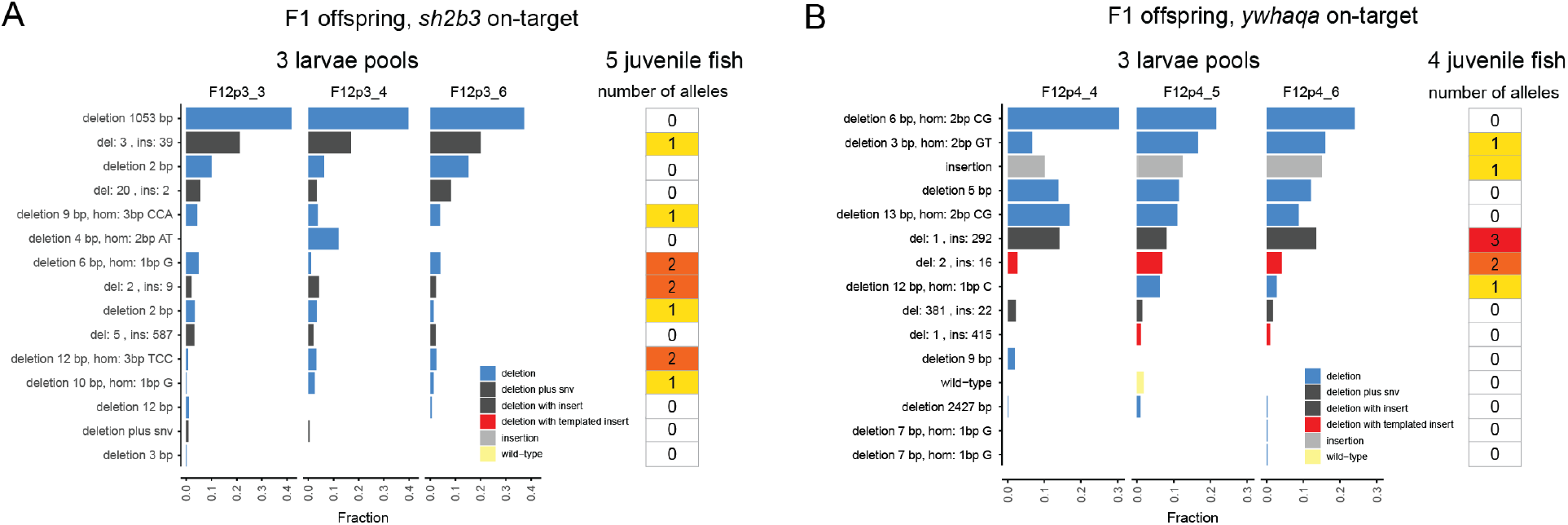
Overview of CRISPR-Cas9 induced on-target events in F1 offspring from the same founder pair. **A)** SIQ image with allele frequencies for the three pools of sh2b3-edited F1 larvae displayed as horizontal bars. The top 15 most frequent events in the F1 larvae are shown in the plot. The column at the right shows the number of occurrences of the editing events among five juvenile fish. **B)** SIQ image with allele frequencies for the three pools of ywhaqa-edited F1 larvae displayed as horizontal bars (at the left) and alleles from four individual juvenile F1 zebrafish (at the right). The 15 most frequent events in the ywhaqa F1 larvae pools are shown in the plot.

### Unintended genome editing outcomes are detectable in the F1 generation

We next analyzed “unintended” outcomes of CRISPR-Cas9 genome editing in the F1 zebrafish. For this reason, we focused on large on-target structural variants and off-target mutations previously detected in the LR-PCR data (4) and aimed to verify their presence in the PureTarget data. In all three samples of pooled *sh2b3*-edited F1 larvae, a deletion of 1053 bp was detected, an event that is also clearly visible in IGV (Figure 5A). This large deletion was the most frequent event in the SIQ analysis of F1 larvae (Figure 4A) but it was not detected in any of the juvenile *sh2b3*-edited F1 zebrafish. One example of off-target genome editing was detected in *ywhaqa* off-target 2. This off-target deletion, which spanned 3bp and was located in an exon of *ywhaqb,* was found to be homozygous in one juvenile fish and heterozygous in two other juveniles (Figure 5B). The same 3 bp deletion in *ywhaqb* was present in all three pools of *ywhaqa*-edited F1 larvae. Our examples thus show that unexpected CRISPR-Cas9 editing outcomes, such as large on-and off- target deletions, can transfer to the next generation. As mentioned above, these examples were earlier using LR-PCR but we here corroborate the findings with PureTarget as an alternative sequencing method.

**Figure 5.**
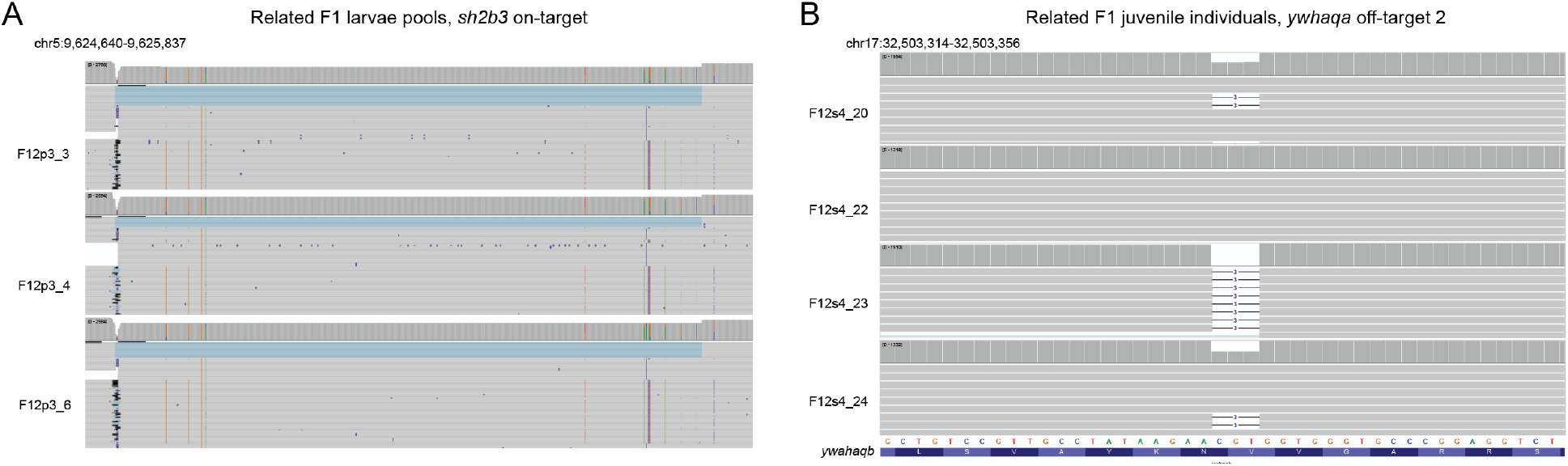
Examples of unintended genome editing events in the F1 generation. **A)** IGV view of a 1053 bp deletion detected in pooled sh2b3-edited F1 larvae. The large deletion is represented by the light blue region that is present in all three samples of pooled larvae. **B)** A three base pair deletion in the ywhaqa off-target 2 region, located inside an exon of the ywhaqa paralogue ywhaqb. The IGV view shows the presence of this off-target mutation with separate tracks for four juvenile F1 zebrafish. Two of the F1 fish are heterozygous, one F1 fish is homozygous for the off-target deletion, and one F1 fish is homozygous for the reference allele.

### CRISPR-Cas9 genome editing has no visible effect on 5mC CpG methylation

PureTarget is an amplification-free protocol for PacBio sequencing. This enables detection of CpG 5mC methylation signals in individual reads and quantification of allele-specific methylation(29). We therefore aimed to investigate whether CRISPR-Cas9 genome editing had a noticeable effect on 5mC methylation signals in our PureTarget experiment. Hence, we compared the methylation patterns in pooled CRISPR-Cas9 edited founder larvae to the signals in the control sample. This analysis was performed only for the on-target regions where CRISPR-Cas9 editing was abundant. By visualizing the data in IGV, we could clearly see that the *sh2b3* on-target region was methylated in the edited founder larvae, while the *ywhaqa* on-target region was unmethylated (Figure 6). Both for *sh2b3* and *ywhaqa*, the methylation patterns were nearly identical in the control sample. Similar results were obtained when examining 5mC methylation signals in the *ldrla* and *nbeal2* on-target regions (supplementary Figure S5). We therefore conclude that CRISPR-Cas9 genome editing has no significant effect on 5mC methylation patterns in our zebrafish experiment. This may be unsurprising since the Cas9 cleavage sites were located in coding exons. However, it is plausible that alterations in methylation could occur in other experiments, i.e. when performing CRISPR-editing of promoter regions or regulatory elements. With PureTarget it would be possible to link the methylation pattern to a genome editing outcome on a single molecule level. In addition, this approach can be used to assess the outcomes of DNA methylation editing(30).

**Figure 6.**
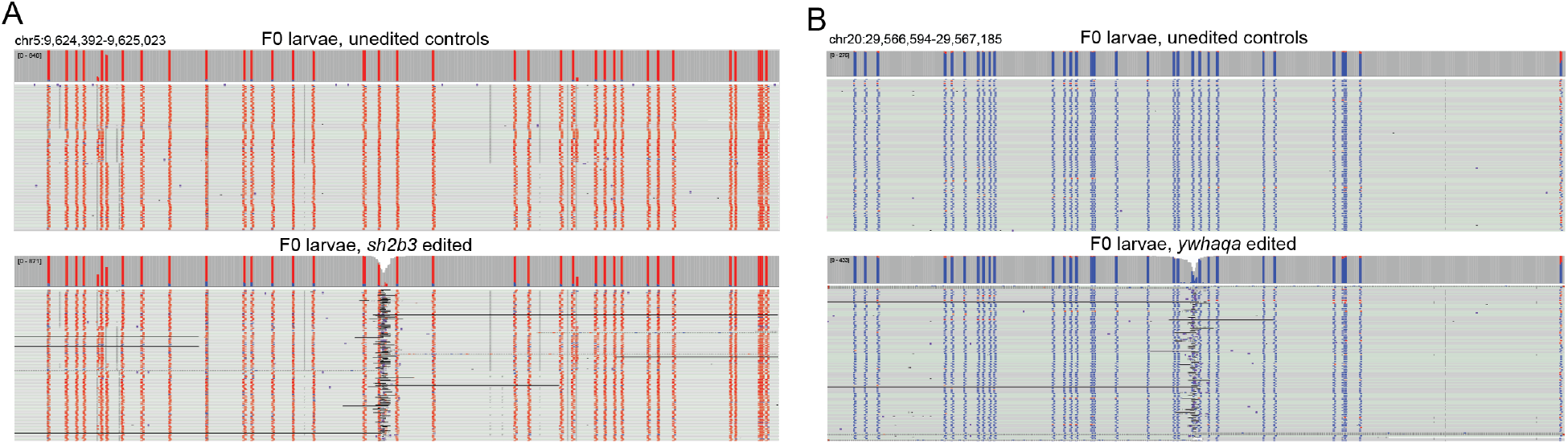
Comparison of 5mC CpG methylation signals between CRISPR-Cas9 edited and control samples. **A)** IGV view showing 5mC signals in the PureTarget data from a sample of pooled sh2b3 founder larvae (bottom) compared with unedited controls (top). The red lines indicate that the CpG signals in this region are methylated both in the sh2b3-edited larvae and unedited controls. **B)** Methylation signals in pooled larvae edited at ywhaqa (bottom) compared with unedited controls (top). The blue lines indicate that the CpG signals in this region are unmethylated both in the edited larvae and controls.

## Discussion

PureTarget generated an average coverage of 1168x for the 30 successfully sequenced samples across seven on-target and off-target sites, corresponding to an enrichment rate of >4000x. This can be compared with the 4.6x enrichment rate previously reported for ONT adaptive sampling(19). Furthermore, PureTarget produced extremely accurate kilobase-length reads, with every read originating from a single molecule. These features make PureTarget an ideal method for verifying CRISPR-Cas9 induced small indels as well as larger SVs, allowing detection of genetic mosaicism down to allele frequencies of 1%. However, PureTarget is based on designing two gRNAs flanking each target to cut out the ROIs. Hence, the method cannot identify events if one of the gRNA binding sites is disrupted, such as >5 kb SVs or larger chromosomal aberrations. In order to capture all possible editing outcomes in a sample, PureTarget might therefore need to be complemented with shallower whole genome sequencing.

While LR-PCR often preferentially amplifies one of the alleles in individual F1 fish, PureTarget gives an even representation of the two haplotypes. This bias in LR-PCR data could be due to length differences between the two haplotypes, or other factors inhibiting the amplification of one of the alleles, like high GC-content or presence of long repeats. To some extent, the LR-PCR results could have been improved by adding UMIs, which was not done in this comparison. However, UMI-based approaches are not perfect either, since errors may occur in the tag sequence(31). Moreover, the UMIs can only remove duplicate molecules, but not other types of amplification biases. Therefore, we believe that an amplification-free enrichment protocol is better suited for analysis of genetic mosaicism. Although ONT’s Cas9 enrichment protocol could be an alternative for generating deep coverage over the target regions(24,25), it lacks the ability of PureTarget to generate extremely high-quality reads. In this study, the single molecule reads reached a QV score of 39, corresponding to one error in approximately 8000 bases.

The PureTarget data gives an unbiased view of the extent of mosaicism generated by CRISPR-Cas9 genome editing. In this study, the CRISPR-Cas9 complex was injected into the fertilized zebrafish eggs at the single cell stage. Our finding of up to 18 unique on-target editing events in a single founder zebrafish implies that CRISPR-Cas9 editing continues to take place after several cell divisions, as described elsewhere(32). Interestingly, mosaicism was also present in the germ cells of founder zebrafish, based on the number of distinct editing events (n=7 and n=5) observed in DNA from pooled F1 zebrafish larvae and juvenile F1 zebrafish obtained from a founder pair. Our finding of widespread mosaicism in the germ cells of founder zebrafish is relevant for breeding programs where CRISPR-Cas9 is used. The results also raise further concerns for germline editing since it is difficult to predict which edits will segregate into future generations. However, it is possible that mosaicism can be mitigated for example by altering the CRISPR-Cas9 conditions, using other delivery systems, or more precise genome editing approaches like base editing(33).

In this experiment, 32 samples were multiplexed on a single Revio SMRTcell. This resulted in a relatively low cost per sample as compared with other amplification-free long-read sequencing methods. Given the large number of reads obtained, it is likely possible to further increase multiplexing and performance, e.g. by improving the target yield and accuracy of the HiFi reads. Enrichment may be optimized by altering the Cas9 concentrations used in the PureTarget protocol and throughput may be increased by modifying the conditions of loading the library on the Revio SMRTcell. Accuracy may be improved by increasing the movie acquisition time of the sequencing run. In this experiment the acquisition time was 24h but this could be increased to 30h to allow the molecules to be sequenced with even more passes. In addition to these methodological changes, perhaps the most important improvements will come from further developments in sequencing chemistry and long-read instrumentation.

PureTarget gives a more detailed view than ever before into CRISPR-Cas9 induced genetic mosaicism. Our results suggest that the method may be useful also for other types of studies where it is important to analyze low-frequency mutations occurring at selected target sites, like in cancer research. Although we did not see any effects of gene editing on methylation in the on-target regions, the feature of detecting 5mC patterns for individual DNA molecules can be highly relevant for example when studying the effects of editing in regulatory regions, or when assessing outcomes of DNA methylation editing(30). Soon it might be possible to extend this analysis to additional base modifications such as 5hmC(34). In summary, the ability to generate over 1000x target coverage of near-perfect native DNA reads opens up for a wide range of applications where it is necessary to study the full extent of genetic variation at specific genomic loci.

## Methods

### Zebrafish handling and CRISPR-Cas9 genome editing

The CRISPR-Cas9 genome editing of zebrafish was performed for an earlier study by Höijer et al(4). Briefly, all zebrafish experiments and husbandry were conducted in accordance with Swedish and European regulations and have been approved by the Uppsala University Ethical Committee for Animal Research (Dnr 5.8.18-13680/2020). The genes of interest were targeted using one gRNA per orthologue with an anticipated editing efficiency >90%. RNA duplexes of the chemically synthesized Alt-R^®^ crRNA (IDT) and Alt-R® tracrRNA (IDT) were complexed with Alt-R® S.p. Cas9 nuclease, v.3 (IDT) to form “duplex guide RNPs” (dgRNPs), as described by Hoshijima K et al(32). The dgRNPs were then injected into fertilized zebrafish eggs at the 1-cell stage. Uninjected embryos were kept and used as controls. The injected founder embryos were raised to adulthood at which time randomly selected mating pairs were in-crossed. Pools of 25–30 10- day-old F0 larvae, individual F0 adult fish, pools of 30 5-day-old F1 larvae, and fin clips of individual juvenile F1 fish were collected throughout the experiment for downstream analyses. Adult zebrafish were sacrificed by prolonged exposure to tricaine, followed by snap freezing in liquid nitrogen to ensure DNA integrity.

### Extraction of genomic DNA from zebrafish

All samples were extracted using the MagAttract HMW DNA Kit (Qiagen) and the “Manual Purification of High-Molecular-Weight Genomic DNA from Fresh or Frozen Tissue” protocol according to the manufacturer’s instructions. A tissue homogenization step using a pestle was added to the protocol prior to the lysis step. DNA integrity of the extracted samples was assessed using the Femto Pulse system (Agilent Technologies) using the Genomic DNA 165 kb kit.

### Design of gRNAs for PureTarget assay

The off-target sites studied in this paper were originally detected by the Nano-OTS method(35). Guide RNAs (gRNAs) for the seven on-targets and off-targets were designed using CHOPCHOP v.3(36) and were ordered as Alt-R™ CRISPR-Cas9 sgRNA from Integrated DNA Technologies (IDT). The size of the enriched targets was designed to be approximately 5 kb in size. Upon arrival the gRNAs were pooled equimolarly to 5 μM and stored for future usage. Further information about the gRNAs is available in Supplementary Table S1.

### PureTarget enrichment of on and off-target sites and PacBio sequencing

1.5 μg of genomic DNA from each of the 32 zebrafish samples was initially processed in a gDNA repair step using a R&D protocol provided by PacBio. In this step, 5 μl of repair buffer (PacBio) and 1 μl of DNA repair mix (PacBio) were added to each sample and incubated at 37°C for 60 min, followed by 52°C for 30 min. Subsequently, the samples were bead-washed using 1x of SMRTbell cleanup beads (PacBio). Following gDNA repair, the 32 samples were processed using the PureTarget repeat expansion panel kit and the “Generating PureTarget repeat expansion panel libraries” protocol (PacBio) according to manufacturer’s instructions. We used our own gRNA pool instead of the repeat expansion panel provided with the kit. All 32 samples were then sequenced on a Revio SMRTcell using the SPRQ chemistry and a 24 hour movie time. SMRT Link v. 25.2 was used for sequencing and data processing.

### Visualizing aligned reads in IGV

To visualize the alignments over the target region, the reads were aligned to the zebrafish reference genome (GRCz11) using the “spliced” parameter in minimap2 v.2.26(37). This parameter is mainly intended for alignment of long RNA data, but we found it to improve alignment for large deletions in the target regions, both for the PureTarget and LR-PCR data. In parallel, another alignment to the GRCz11 reference genome was performed in SMRTLink v25.2 for the PureTarget data, using default parameters. This yielded alignment files that include 5mC methylation signals. The aligned data was visualized in IGV v 2.19.4(27).

### PureTarget coverage analysis and quality assessment

Two of the zebrafish samples had less than 30X coverage for all target regions and were removed from further analysis. For the remaining 30 samples, the average coverage across all sites was obtained by applying the ‘coverage’ function in samtools v1.20(38) to the data aligned to the regions of interest. To calculate quality scores, fastq files were extracted for each of the sites using samtools ‘bam2fq’ and FastQC v0.11.9 (*www.bioinformatics.babraham.ac.uk/projects/fastqc*)was used to generate phred quality score distributions for each sample and target.

### Calculating the PureTarget enrichment rate

The GRCz11 chromosomal sequences consist of 1.345 Gb and the combined size of the seven target regions in the PureTarget design is 33,258 bp. This means that the targets cover about 0.0025% of the zebrafish reference genome. In total, the PureTarget sequencing generated 10.4 Gb HiFi data from one Revio SMRTcell. If sequencing was completely random, we expect a mean coverage of 0.243x for each of the 32 samples over target regions. However, in the PureTarget experiment we reach a mean target coverage of 1096x for the 32 samples across the seven on- and off-target regions. This implies that PureTarget has an enrichment rate of about 4500x as compared with shotgun whole genome sequencing. When removing the two samples that generated less than 30x coverage for all target regions, the mean target coverage was 1168x. This corresponds to an enrichment rate of 4800x.

### Detection of and visualization of genome editing events using SIQ

Analysis of CRISPR-Cas9 genome editing outcomes was performed using SIQ v2.1 (28) and the results were visualized using SIQplotteR (28) As input to SIQ, we used fastq files extracted from each mapped bam file for each sample/target combination. The analysis was run both for the PureTarget and LR-PCR data. Only events occurring within 2bp of the expected CRISPR-Cas9 genome editing site were considered in the SIQ analysis, and a cut-off of 1% was applied for the detection of mosaic events in a sample.

## Acknowledgements

We want to thank Sarah Kingan and Nadia Sellami at PacBio for their assistance in designing the project. Library preparation was done at the SciLifeLab National Genomics Infrastructure (NGI) in Uppsala, Sweden and PacBio sequencing was performed using instrumentation at Clinical Genomics Uppsala (CGU). Computational resources were provided by the National Academic Infrastructure for Supercomputing in Sweden (NAISS) at UPPMAX. The work was funded by a technology development project grant from SciLifeLab NGI. Some of the reagents were provided free of charge by PacBio.

## Notes

### Competing Interest Statement

Some of the sequencing reagents used in this study were provided free of charge by PacBio.

