## Supplementary Figures and Tables for "Accurate characterization of CRISPR-Cas9 genome editing outcomes and mosaicism with near-perfect long reads"

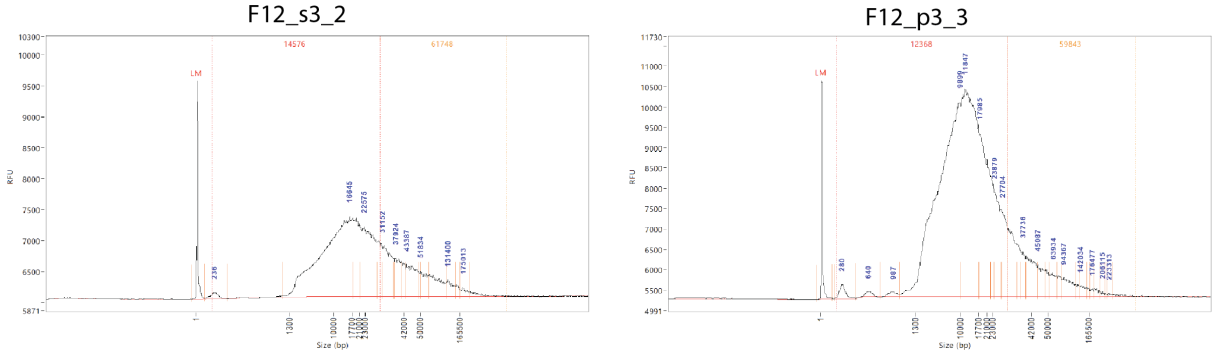


**Supplementary Figure S1. Quality of zebrafish DNA.** Examples of DNA integrity of the zebrafish DNA samples. The DNA integrity was assessed using the Agilent FemtoPulse system and the Genomic DNA 165 kb kit. The sample to the left is DNA extracted from an individual zebrafish and the sample to the right is DNA extracted from a pool of zebrafish larvae.

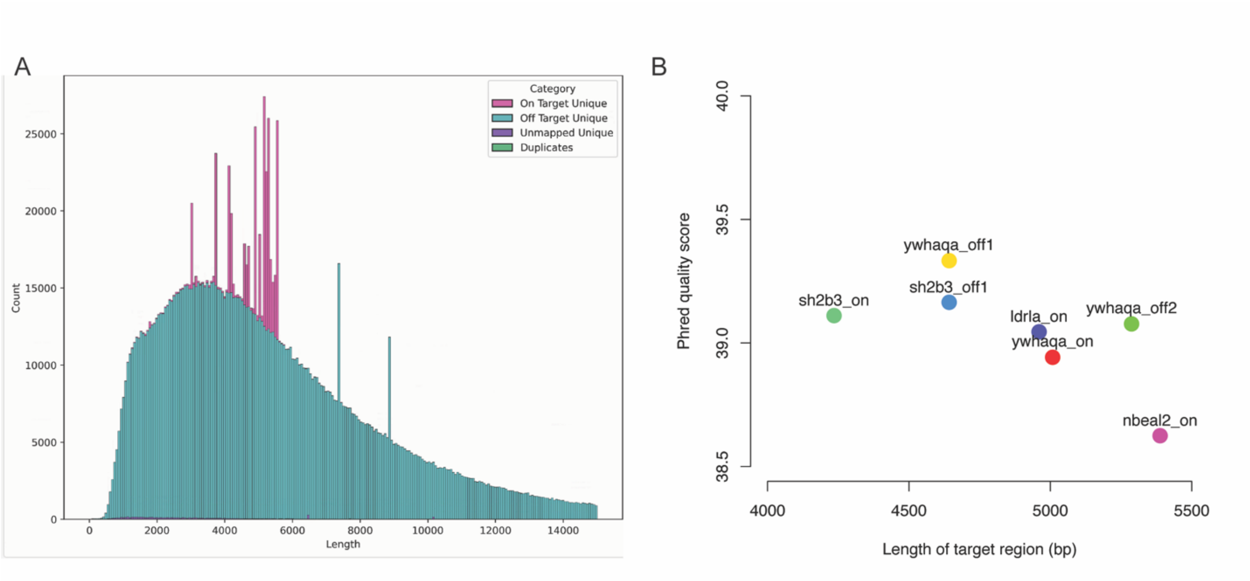


**Supplementary Figure S2. Read length and quality of PureTarget sequencing. A)** Read length distribution of the PureTarget sequencing run. The pink bars correspond to reads that map to the target regions. **B)** Distribution of quality scores for PureTarget on-target reads. On the y-axis are the average phred quality scores for all reads mapping to the seven targets in the PureTarget panel. All seven targets have phred scores between 38.5 and 39.5, with slightly higher scores for shorter target regions.

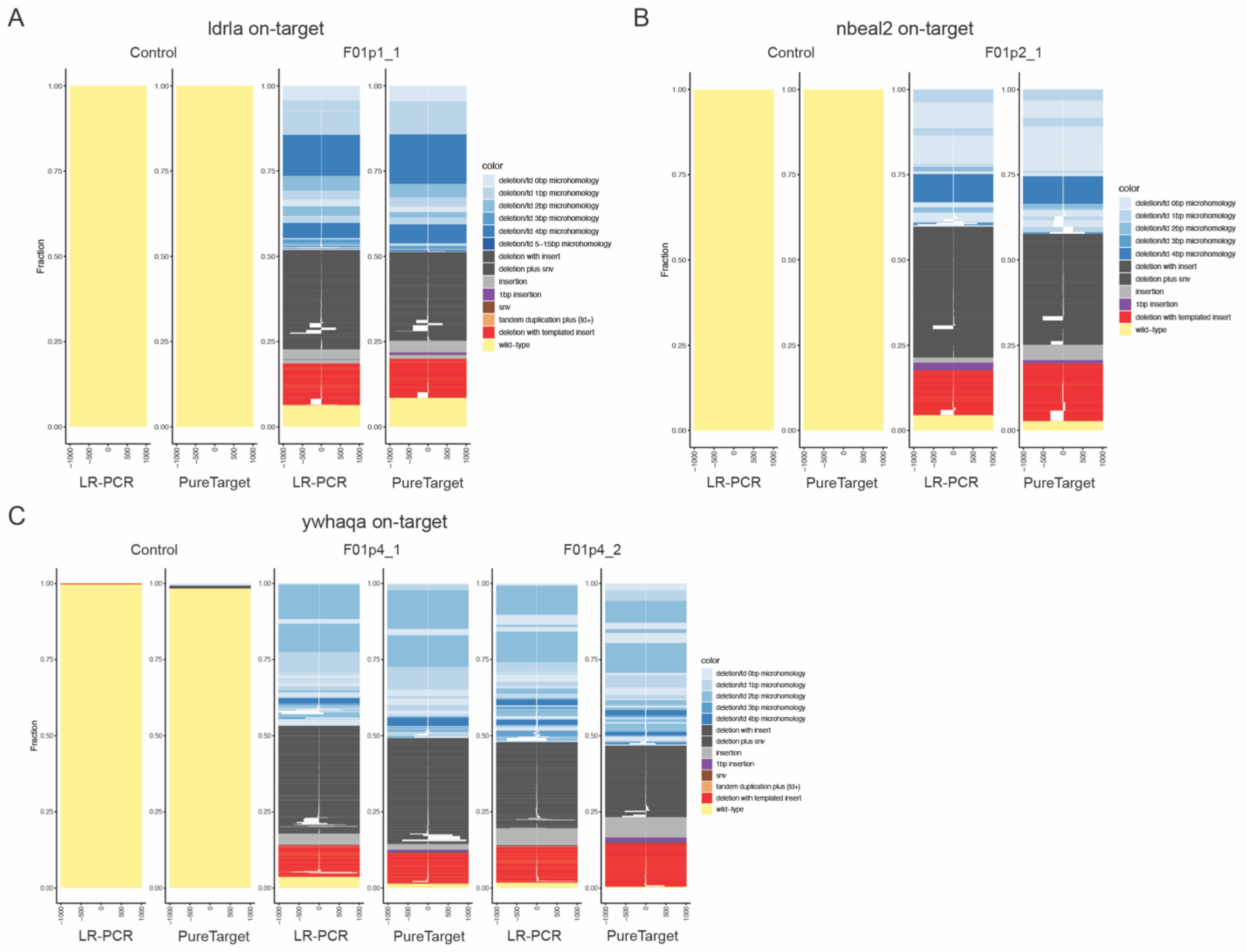


**Supplementary Figure S3. CRISPR-Cas9 induced genetic mosaicism at on-target sites for pools of founder zebrafish larvae. A)** SIQ tornado plots showing the distribution of on-target genome editing outcomes in the uninjected control sample and a pool of *ldrla* founder (F0) larvae. For each sample, outcomes are shown both for LR-PCR and PureTarget data in a window centered around the CRISPR-Cas9 editing site. **B)** SIQ tornado plots for an uninjected control sample and a pool of *nbeal2* founder (F0) larvae. **C)** SIQ tornado plots for an uninjected control sample and two pools of *ywhaqa* founder (F0) larvae.

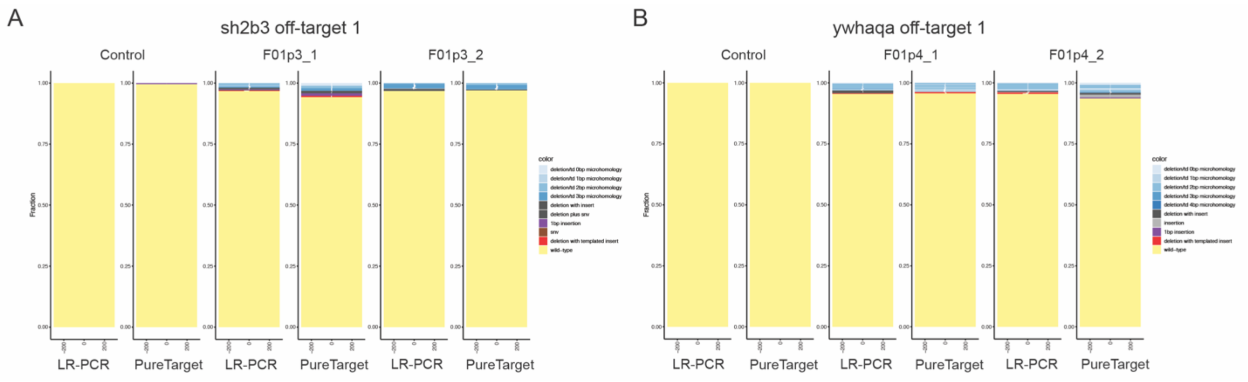


**Supplementary Figure S4. CRISPR-Cas9 induced genetic mosaicism at off-target sites for pools of founder zebrafish larvae. A)** SIQ tornado plots showing the distribution of off-target genome editing outcomes at *sh2b3* off-target site 1 in the uninjected control sample and two pools of *sh2b3* founder (F0) larvae. For each sample, outcomes are shown both for LR-PCR and PureTarget data in a window centered around the CRISPR-Cas9 editing site. **B)** SIQ tornado plots showing the distribution of off-target genome editing outcomes for an uninjected control sample and two pools of *ywhaqa* founder (F0) larvae at *ywhaqa* off-target site 1.

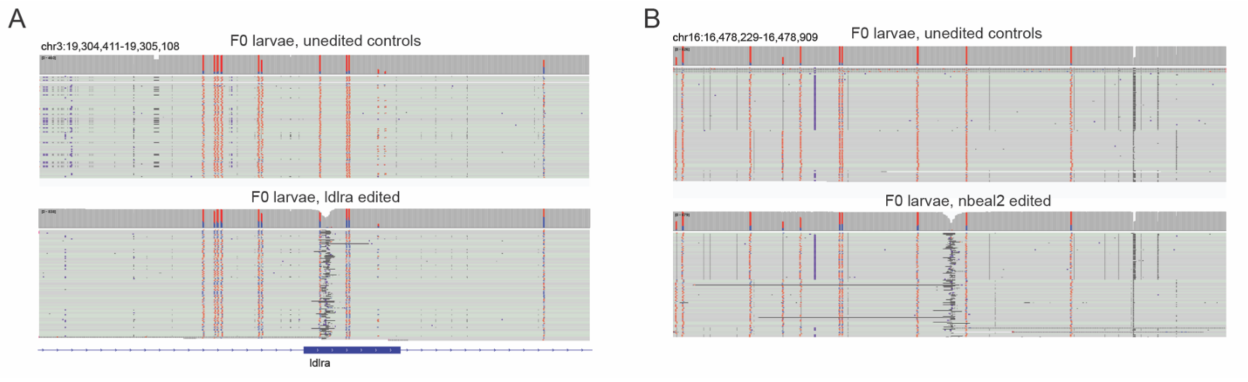


**Supplementary Figure S5. Comparison of 5mC CpG methylation signals between CRISPR-Cas9 edited and control samples. A)** IGV view showing 5mC signals in the PureTarget data from a pool of *ldlra* founder larvae (bottom) compared with a sample of pooled unedited controls (top). **B)** Methylation signals in a pool of *nbeal2* founder larvae (bottom) compared with a sample of pooled controls (top).

### Supplementary Tables

#### Supplementary Table S1. Information on PureTarget gRNAs and regions of interest (ROIs)

| Name | Strand | Sequence | PAM | Genomic coordinates | ROI size (bp) |
|---|---|---|---|---|---|
| ldlra_on_fwd | + | TAATAATTTTCTACGTCCTA | CGG | chr3:19,302,176-19,302,195 | 4961 |
| ldlra_on_rev | - | GTCAACATAGAAATTTCAGA | TGG | chr3:19,307,117-19,307,136 | |
| nbeal2_on_fwd | + | AGAGCTGAAAGGTAAGGACT | GGG | chr16:16,475,683-16,475,702 | 5389 |
| nbeal2_on_rev | - | TAGTTCATGACCTAGACCTC | AGG | chr16:16,481,052-16,481,071 | |
| sh2b3_off1_fwd | + | AACCACATAACCTTTTCTCC | CGG | chr15:31,595,678-31,595,697 | 4643 |
| sh2b3_off1_rev | - | AATTCCTCAATTCTAGGATA | GGG | chr15:31,600,301-31,600,320 | |
| sh2b3_on_fwd | + | TTAAAGCCAACTGGTTCCCA | CGG | chr5:9,622,006-9,622,025 | 4236 |
| sh2b3_on_rev | - | GATATTCACCACACCTCAGT | TGG | chr5:9,626,222-9,626,241 | |
| ywhaqa_off1_fwd | + | CGGATTGGGCAAATGCCAGA | AGG | chr5:3,542,442-3,542,461 | 3739 |
| ywhaqa_off1_rev | - | ATGGTAACCACGCACTGTTT | TGG | chr5:3,546,161-3,546,180 | |
| ywhaqa_off2_fwd | + | CTCAAAGGCTATCAATCCGG | TGG | chr17:32,500,471-32,500,490 | 5288 |
| ywhaqa_off2_rev | - | ACTAGAGTGCTGACAATGAT | GGG | chr17:32,505,739-32,505,758 | |
| ywhaqa_on_fwd | + | TAGATAAGCCAAGTAGACTC | TGG | chr20:29,564,222-29,564,241 | 5009 |
| ywhaqa_on_rev | - | CTACTTTAAAGGTGCCCAGA | AGG | chr20:29,569,211-29,569,230 | |

#### Supplementary Table S2. Table with all 32 zebrafish samples run with PureTarget

| Sample name | Generation | Edit | Age | Mating pair | Sample ID |
|---|---|---|---|---|---|
| F12p3_3 | F1 | *sh2b3* | 5 dpf 30 larvae | sh2b3 pair 3 | pr_180_001 |
| F12p3_4 | F1 | *sh2b3* | 5 dpf 30 larvae | sh2b3 pair 3 | pr_180_002 |
| F12p3_6 | F1 | *sh2b3* | 5 dpf 30 larvae | sh2b3 pair 3 | pr_180_003 |
| F12s3_2 | F1 | *sh2b3* | Juvenile fin | sh2b3 pair 3 | pr_180_004 |
| F12s3_3 | F1 | *sh2b3* | Juvenile fin | sh2b3 pair 3 | pr_180_005 |
| F12s3_12 | F1 | *sh2b3* | Juvenile fin | sh2b3 pair 3 | pr_180_006 |
| F12s3_13 | F1 | *sh2b3* | Juvenile fin | sh2b3 pair 3 | pr_180_007 |
| F12s3_14 | F1 | *sh2b3* | Juvenile fin | sh2b3 pair 3 | pr_180_008 |
| F12p4_4 | F1 | *ywhaqa* | 5 dpf 30 larvae | ywhaqa pair 2 | pr_180_009 |
| F12p4_5 | F1 | *ywhaqa* | 5 dpf 30 larvae | ywhaqa pair 2 | pr_180_010 |
| F12p4_6 | F1 | *ywhaqa* | 5 dpf 30 larvae | ywhaqa pair 2 | pr_180_011 |
| F12s4_20 | F1 | *ywhaqa* | Juvenile fin | ywhaqa pair 2 | pr_180_012 |
| F12s4_22 | F1 | *ywhaqa* | Juvenile fin | ywhaqa pair 2 | pr_180_013 |
| F12s4_23 | F1 | *ywhaqa* | Juvenile fin | ywhaqa pair 2 | pr_180_014 |
| F12s4_24 | F1 | *ywhaqa* | Juvenile fin | ywhaqa pair 2 | pr_180_015 |
| F01wt_1 | F0 | *wt* | 10 dpf 30 larvae | | pr_180_016 |

| | | | | | |
|---|---|---|---|---|---|
| F01p3_1 | F0 | *sh2b3* | 10 dpf 30 larvae | | pr_180_017 |
| F01p3_2 | F0 | *sh2b3* | 10 dpf 30 larvae | | pr_180_018 |
| F01p4_1 | F0 | *ywhaqa* | 10 dpf 30 larvae | | pr_180_019 |
| F01p4_2 | F0 | *ywhaqa* | 10 dpf 30 larvae | | pr_180_020 |
| F01p1_1 | F0 | *ldlra* | 10 dpf 30 larvae | | pr_180_021 |
| F01p2_1 | F0 | *nbeal2* | 10 dpf 25 larvae | | pr_180_022 |
| F01s3_2 | F0 | *sh2b3* | adult | | pr_180_023 |
| F01s3_3* | F0 | *sh2b3* | adult | | pr_180_024 |
| F01s3_4 | F0 | *sh2b3* | adult | | pr_180_025 |
| F01s3_5* | F0 | *sh2b3* | adult | | pr_180_026 |
| F01s3_6 | F0 | *sh2b3* | adult | | pr_180_027 |
| F01s4_3 | F0 | *ywhaqa* | adult | | pr_180_028 |
| F01s4_4 | F0 | *ywhaqa* | adult | | pr_180_029 |
| F01s4_6 | F0 | *ywhaqa* | adult | | pr_180_030 |
| F01s4_15 | F0 | *ywhaqa* | adult | | pr_180_031 |
| F01s4_16 | F0 | *ywhaqa* | adult | | pr_180_032 |

*These two samples failed to generate 30X coverage over the target sites and were excluded from the analysis

**Supplementary Table S3. PureTarget run statistics for one Revio 8M SMRT cell**

| | |
|---|---|
| HiFi Bases | 10,377,051,654 |
| HiFi Reads | 1,826,314 |
| Median HiFi Read Length (bp) | 4,925 |
| Median HiFi Read Quality | QV41 |
| Sample count | 32 |
| Target regions | 7 |
| Total Target Length (bp) | 33,258 |
| Percent of bases mapping to target regions (min-max for the 32 samples) | 7.1% - 18.4% |

*Results obtained by running the Target Enrichment plugin in SMRTLink v.25.2

#### Supplementary Table S4. Number of PureTarget reads mapping to the seven target sites

| Sample | *ywhaqa* on1 | *ywhaqa* off1 | *ywhaqa* off2 | *sh2b3* on | *sh2b3* off1 | *ldrla* on | *nbeal2* on | all target reads | total reads | % on target |
|---|---|---|---|---|---|---|---|---|---|---|
| F12p3_3 | 1314 | 1842 | 2686 | 4602 | 1416 | 1238 | 2138 | 15236 | 83948 | 18.1 |
| F12p3_4 | 1364 | 1812 | 2940 | 4996 | 1376 | 1148 | 2302 | 15938 | 79722 | 20 |
| F12p3_6 | 1548 | 1954 | 2980 | 4844 | 1472 | 1482 | 2216 | 16496 | 74211 | 22.2 |
| F12s3_2 | 698 | 1326 | 1204 | 3146 | 860 | 908 | 1110 | 9252 | 82765 | 11.2 |
| F12s3_3 | 448 | 756 | 940 | 2640 | 614 | 698 | 880 | 6976 | 72641 | 9.6 |
| F12s3_12 | 880 | 1444 | 1978 | 3138 | 858 | 988 | 1780 | 11066 | 55056 | 20.1 |
| F12s3_13 | 716 | 1250 | 1446 | 3400 | 698 | 804 | 1212 | 9526 | 47696 | 20 |
| F12s3_14 | 666 | 736 | 1168 | 2704 | 658 | 432 | 1056 | 7420 | 34783 | 21.3 |
| F12p4_4 | 1364 | 1928 | 2904 | 4814 | 1434 | 1802 | 2528 | 16774 | 57986 | 28.9 |
| F12p4_5 | 1286 | 1690 | 2354 | 3826 | 1252 | 1638 | 2204 | 14250 | 43903 | 32.5 |
| F12p4_6 | 1128 | 1784 | 2650 | 4012 | 1262 | 1548 | 2122 | 14506 | 46441 | 31.2 |
| F12s4_20 | 584 | 946 | 1566 | 3020 | 618 | 976 | 1302 | 9012 | 40312 | 22.4 |
| F12s4_22 | 460 | 818 | 1222 | 1648 | 406 | 802 | 992 | 6348 | 19183 | 33.1 |
| F12s4_23 | 1224 | 2310 | 2032 | 5942 | 1218 | 1968 | 2288 | 16982 | 30532 | 55.6 |
| F12s4_24 | 616 | 1166 | 1346 | 2284 | 666 | 778 | 1152 | 8008 | 27441 | 29.2 |
| F01wt_1 | 560 | 1036 | 1606 | 1700 | 764 | 950 | 1278 | 7894 | 49305 | 16 |
| F01p3_1 | 676 | 1634 | 1736 | 3034 | 846 | 1268 | 1584 | 10778 | 100440 | 10.7 |
| F01p3_2 | 842 | 2122 | 2142 | 4014 | 1166 | 1602 | 2028 | 13916 | 122110 | 11.4 |
| F01p4_1 | 956 | 1994 | 2328 | 3536 | 1156 | 1510 | 1964 | 13444 | 81800 | 16.4 |
| F01p4_2 | 630 | 1452 | 1786 | 3030 | 804 | 1024 | 1584 | 10310 | 72999 | 14.1 |
| F01p1_1 | 978 | 1738 | 2658 | 3928 | 1196 | 1760 | 2250 | 14508 | 71502 | 20.3 |
| F01p2_1 | 572 | 1202 | 1492 | 2526 | 796 | 1234 | 1472 | 9294 | 60767 | 15.3 |
| F01s3_2 | 300 | 930 | 622 | 1556 | 420 | 500 | 552 | 4880 | 56562 | 8.6 |
| F01s3_3* | 2 | 2 | 4 | 2 | 0 | 0 | 2 | 12 | 5633 | 0.2 |
| F01s3_4 | 70 | 224 | 150 | 176 | 108 | 124 | 148 | 1000 | 31796 | 3.1 |
| F01s3_5* | 6 | 64 | 16 | 14 | 8 | 8 | 4 | 120 | 20914 | 0.6 |
| F01s3_6 | 378 | 928 | 554 | 1002 | 378 | 474 | 466 | 4180 | 60724 | 6.9 |
| F01s4_3 | 940 | 2696 | 3166 | 5888 | 156 | 2090 | 2602 | 17538 | 62493 | 28.1 |
| F01s4_4 | 286 | 624 | 624 | 838 | 22 | 470 | 412 | 3276 | 25010 | 13.1 |
| F01s4_6 | 1020 | 1616 | 2402 | 3862 | 1234 | 1744 | 2040 | 13918 | 110901 | 12.5 |
| F01s4_15 | 404 | 570 | 732 | 604 | 480 | 454 | 664 | 3908 | 48161 | 8.1 |
| F01s4_16 | 216 | 754 | 606 | 746 | 142 | 370 | 426 | 3260 | 38142 | 8.5 |
| | | | | | | | | 310026 | 1815879 | 17.1 |

*These two samples failed to generate 30X coverage over the target sites and were excluded from the analysis

**Supplementary Table S5. Allelic distribution of reads at heterozygous SNVs in F1 fish**

| Sample | Method | Target | SNV position* (ref/alt) | Total | Ref | Alt |
|---|---|---|---|---|---|---|
| F12s3_2 | Amplicon | sh2b3 on | chr5:9624430 (C/A) | 1102 | 372 (34%) | 729 (66%) |
| F12s3_2 | PureTarget | sh2b3 on | chr5:9624430 (C/A) | 1582 | 762 (48%) | 820 (52%) |
| F12s3_12 | Amplicon | sh2b3 on | chr5:9624430 (C/A) | 1402 | 177 (13%) | 1225 (87%) |
| F12s3_12 | PureTarget | sh2b3 on | chr5:9624430 (C/A) | 2098 | 1074 (51%) | 1024 (49%) |
| F12s3_13 | Amplicon | sh2b3 on | chr5:9624430 (C/A) | 1396 | 109 (8%) | 1287 (92%) |
| F12s3_13 | PureTarget | sh2b3 on | chr5:9624430 (C/A) | 1702 | 890 (52%) | 812 (48%) |
| F12s4_20 | Amplicon | ywhaqa on | chr20:29566589 (T/C) | 1597 | 1 (0%) | 1595 (100%) |
| F12s4_20 | PureTarget | ywhaqa on | chr20:29566589 (T/C) | 574 | 272 (47%) | 302 (53%) |
| F12s4_20 | Amplicon | ywhaqa off2 | chr17:32502859 (T/C) | 2911 | 282 (10%) | 2629 (90%) |
| F12s4_20 | PureTarget | ywhaqa off2 | chr17:32502859 (T/C) | 1556 | 688 (44%) | 868 (56%) |
| F12s4_22 | Amplicon | ywhaqa on | chr20:29566589 (T/C) | 1567 | 3 (0%) | 1563 (100%) |
| F12s4_22 | PureTarget | ywhaqa on | chr20:29566589 (T/C) | 456 | 234 (51%) | 222 (49%) |
| F12s4_22 | Amplicon | ywhaqa off1 | chr5:3544380 (C/T) | 1539 | 1287 (84%) | 252 (16%) |
| F12s4_22 | PureTarget | ywhaqa off1 | chr5:3544380 (C/T) | 818 | 378 (46%) | 440 (54%) |
| F12s4_22 | Amplicon | ywhaqa off2 | chr17:32502859 (T/C) | 4273 | 479 (11%) | 3793 (89%) |
| F12s4_22 | PureTarget | ywhaqa off2 | chr17:32502859 (T/C) | 1218 | 608 (50%) | 610 (50%) |
| F12s4_23 | Amplicon | ywhaqa on | chr20:29566589 (T/C) | 1164 | 18 (2%) | 1145 (98%) |
| F12s4_23 | PureTarget | ywhaqa on | chr20:29566589 (T/C) | 1212 | 608 (50%) | 604 (50%) |
| F12s4_23 | Amplicon | ywhaqa off1 | chr5:3543062 (C/T) | 1000 | 729 (73%) | 269 (27%) |
| F12s4_23 | PureTarget | ywhaqa off1 | chr5:3543062 (C/T) | 2304 | 1136 (49%) | 1168 (51%) |
| F12s4_24 | Amplicon | ywhaqa off1 | chr5:3543062 (C/T) | 1765 | 1374 (78%) | 389 (22%) |
| F12s4_24 | PureTarget | ywhaqa off1 | chr5:3543062 (C/T) | 1162 | 546 (47%) | 616 (53%) |

*For each amplicon, a representative SNV position was selected for calculating the allele frequency. The SNVs were selected to be located outside copy number variable regions and gaps in the GRCz11 reference.
